# METHimpute: Imputation-guided construction of complete methylomes from WGBS data

**DOI:** 10.1101/190223

**Authors:** Aaron Taudt, David Roquis, Amaryllis Vidalis, René Wardenaar, Frank Johannes, Maria Colomé-Tatché

**Affiliations:** European Research Institute for the Biology of Ageing, University of Groningen, University Medical Centre Groningen, A. Deusinglaan 1, NL-9713 AV Groningen, The Netherlands.; Department of Plant Sciences, Hans Eisenmann-Zentrum for Agricultural Sciences, Technical University Munich, Liesel-Beckmann-Str. 2, 85354 Freising, Germany.; Institute of Computational Biology, Helmholtz Zentrum München, Ingolstädter Landstr. 1, 85764 Neuherberg, Germany.

**Keywords:** methylation, whole-genome bisulfite sequencing, imputation, Hidden Markov Model

## Abstract

Whole-genome Bisulfite sequencing (WGBS) has become the standard method for interrogating plant methylomes at base resolution. However, deep WGBS measurements remain cost prohibitive for large, complex genomes and for population-level studies. As a result, most published plant methylomes are sequenced far below saturation, with a large proportion of cytosines having either missing data or insufficient coverage. Here we present METHimpute, a Hidden Markov Model (HMM) based imputation algorithm for the analysis of WGBS data. Unlike existing methods, METHimpute enables the construction of complete methylomes by inferring the methylation status and level of all cytosines in the genome regardless of coverage. Application of METHimpute to maize, rice and Arabidopsis shows that the algorithm infers cytosine-resolution methylomes with high accuracy from data as low as 6X, compared to data with 60X, thus making it a cost-effective solution for large-scale studies. Although METHimpute has been extensively tested in plants, it should be broadly applicable to other species.

## Introduction

Cytosine methylation (5mC) is a widely conserved epigenetic mark [1–4] with important roles in the regulation of gene expression and the silencing of transposable elements (TEs) and repeats [5,6]. Experimentally-induced changes in 5mC patterns have been shown to affect plant phenotypes [7–9], rates of meiotic recombination [10–13], genome stability [14–18] and alter plant-environment interactions [19–22]. Similar to genetic mutations, changes in 5mC patterns can also occur spontaneously as a result of errors in DNA methylation maintenance [23–26]. There is substantial evidence in plants that experimentally-induced as well as spontaneously occurring 5mC changes can be stably inherited across multiple generations, independently of genetic changes [27]. Cytosine methylation has therefore emerged as a potentially important factor in plant evolution [28–30] and as a possible molecular target for the improvement of commercial crops [31,32].

Plant methylomes are now routinely studied using whole-genome bisulfite sequencing (WGBS), a next generation sequencing (NGS) method that can interrogate the methylation status of individual cytosines at the genome-wide scale. The application of this technology has been instrumental in dissecting the molecular pathways that establish and maintain 5mC patterns in plant genomes. Unlike in animals, plants methylate cytosines in context CG, but also extensively in contexts CHG and CHH, where H = A, T, C [5]. Methylation at CG dinucleotides (mCG) is maintained by methyltransferase 1 (MET1), which is recruited to hemi-methylated CG sites in order to methylate the complimentary strand in a template-dependent manner during DNA replication [33]. By contrast, mCHG is maintained dynamically by the plant specific chromomethylase 3 (CMT3) [34], and requires continuous interactions with H3K9me2 (dimethylation of lysine 9 on histone 3) [35]. Asymmetrical methylation of CHH sites (mCHH) is established and maintained by another member of the CMT family, CMT2 [2,36]. Similar to CMT3, CMT2 dynamically methylates CHH in H3K9me2-associated regions. In addition to these context-specific maintenance mechanisms, all three sequence contexts can also be methylated de novo via RNA-directed DNA methylation (RdDM) [5], which involves short-interfering 24 nucleotide small RNAs (siRNA) that guide the de novo methyltransferase domains rearranged methyltransferase 2 (DRM2) to homologous target sites throughout the genome [37,38].

Although these methylation pathways appear to be broadly conserved across plant species, recent data indicates that there is extensive variation in 5mC patterns both between but also within species [3,39]. Efforts to explore the origin of this variation and its implications for plant evolution, ecology and agriculture will require large inter‐ and intraspecific methylome datasets. Such datasets are currently emerging. To date, the methylomes of over 50 plant species have been analyzed using WGBS [3, 4], including representative species of major taxonomic groups such as angiosperms (flowering plants), gymnosperms, ferns, and non-vascular plants. In addition, the methylomes of over 1000 natural A. thaliana accessions are now available [40], as well as those of several experimentally derived populations [41]. However, deep inter‐ and intraspecific WGBS measurements remain cost-prohibitive, particularly for species with large genomes. Most published plant methylomes have therefore been sequenced far below saturation (i.e. large number of cytosines in the genome are not covered). Indeed, even simple genomes, like that of the model plant *A. thaliana* (Col-0 accession), are typically only sequenced to about 10-30X. At this depth, about 5-10% of cytosines have missing data (*i.e*. zero read coverage) and about 15-20% have nearly uninformative read coverage (< 3 reads), and this problem is exacerbated in more complex genomes, like those of rice and maize (see Fig. 1).

**Figure 1.**
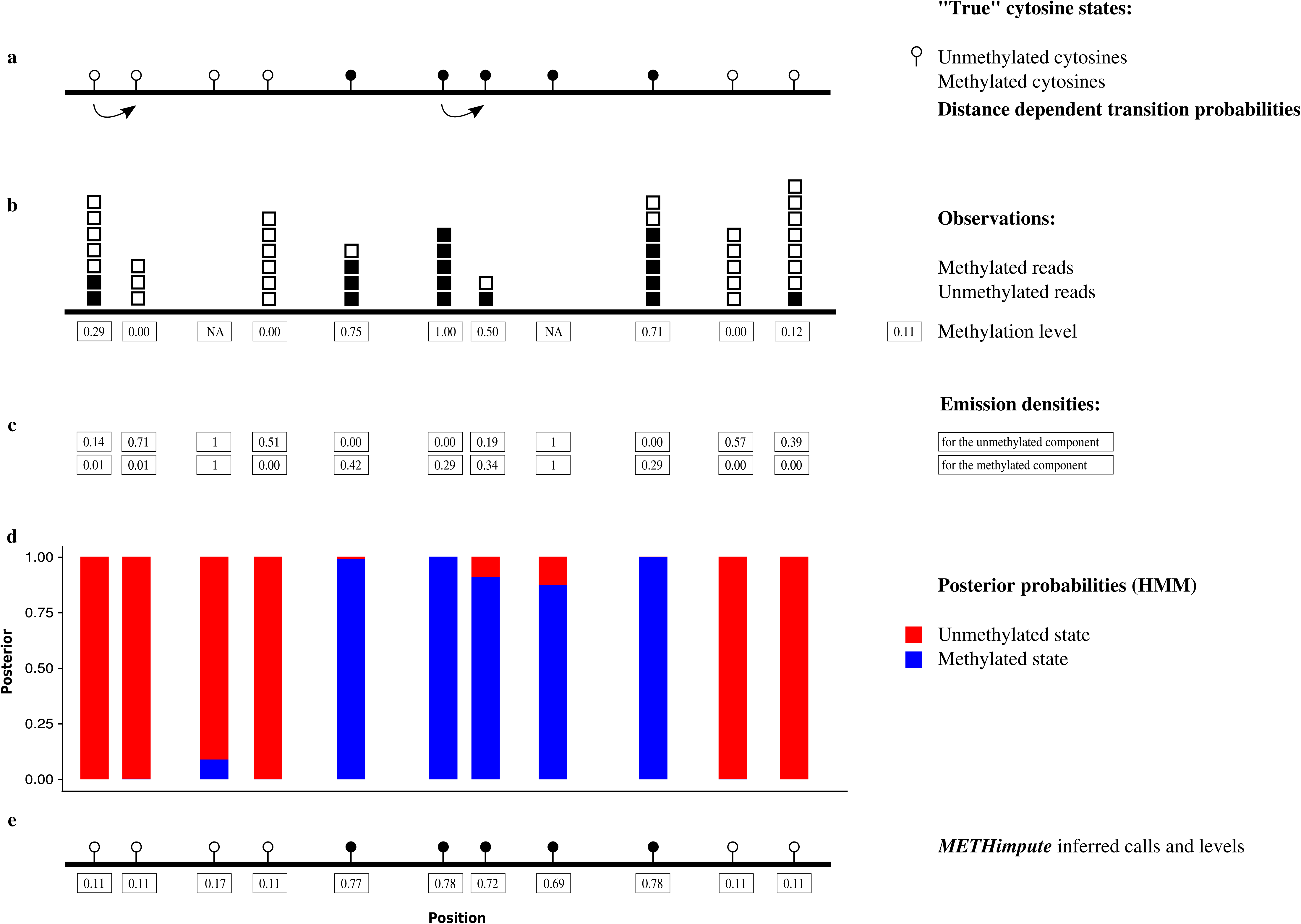
Coverage distributions. **(a-c)** Percentage of cytosines with X coverage (strand-specific). **(d-f)** Percentage of cytosines with missing data (red) and "uninformative” coverage (green), defined as less than three reads.

Low to moderate sequencing depths in individual samples have cumulative consequences for analyzing population-level data. For instance, in the recently released 1000 *A. thaliana* methylome data [40] (measured at 5X coverage per strand on average), 92% of cytosines have missing data in at least one sample when 100 accessions are compared (Fig. SI-1). These incomplete measurements will reduce statistical power in genome-wide methylation QTL (meQTL) mapping studies, in estimates of epimutation rates, or in ecological studies that aim to correlate site-specific methylation levels with environmental/climatic variables. Moreover, incomplete measurements also complicate and potentially bias methylome scans for signature of epigenetic selection using methylation site frequency spectrum (mSFS) analytic approaches [28]. One way to circumvent the missing data problem is to calculate methylation levels over larger regions, ranging anywhere from several hundred to several thousand basepairs and to use these methylation levels for downstream population-level analyses. In the above-mentioned A. thaliana population data, only 36% of 100bp regions in the genome are missing in at least one sample of the 100 accessions, compared with 92% of individual cytosines, and this percentage further decreases with larger region sizes. However, while region-based methylation levels are useful measures for descriptive and correlative analyses, these measures obscure detailed insights into the cytosine-level methylation status calls, and thus arguably undermine the key advantages of WGBS over other lower resolution technologies such as MeDIP-seq. Cytosine-level status calls are needed to be able to apply existing population (epi)genetic models to population methylome data, and to be able to test explicit hypotheses about the evolutionary forces that shape methylome variation patterns within and among species [28].

In order to maximize the information contained in WGBS data and to facilitate cost-effective sequencing decisions for future studies, we developed METHimpute, a Hidden Markov Model (HMM) based imputation algorithm for the construction of base-resolution methylomes from WGBS data. The unique feature of this algorithm is its ability to impute the methylation status and level of cytosines with missing or uninformative measurements, thus yielding complete methylomes even with low-coverage WGBS datasets. Indeed, using published WGBS data from *Arabidopsis thaliana* (rockcress), *Oryza sativa* (rice) and *Zea mays* (maize), we demonstrate that METHimpute accurately reconstructs base-resolution methylomes from data with an average coverage as low as 6X, suggesting that typical sequencing costs could be cut by more than 50% without a significant loss of information.

## Conceptual overview

WGBS is an NGS-based method in which DNA is treated with sodium bisulfite prior to sequencing in order to convert unmethylated cytosines into uracils and finally into thymines during PCR amplification. Hence, a cytosine in a bisulfite treated read that maps to a cytosine in the reference genome provides evidence for methylation, while a thymine that maps to a cytosine does not. Many specialized short read mapping programs make use of this information and output so-called *methylation levels* [42–44]; that is, the proportion of aligned reads that support that a cytosine is methylated out of all the reads covering the site. Methylation levels are inherently noisy due to inefficiencies in the sodium bisulfite conversion step. Moreover, tissue heterogeneity and the highly dynamic maintenance methylation at CHH and CHG, which requires feedback loops with histone modifications and small RNAs [5,6], lead to intermediate methylation levels which are very susceptible to experimental variation. Finally, in WGBS data a large proportion of cytosines are often either not covered by any sequencing read or are covered only by a few number of reads (Fig. 1), meaning that methylation levels at these positions cannot be determined.

To overcome these limitations we developed METHimpute, a Hidden Markov Model (HMM) for the construction of base-resolution methylomes from WGBS data. METHimpute takes methylated and unmethylated read counts at every cytosine as input, and outputs discrete methylation status calls (unmethylated or methylated), together with recalibrated methylation levels between 0 and 1 for every cytosine in the genome, regardless of coverage (Fig. 2).

The METHimpute algorithm fits a two-state HMM to the observed methylation counts. The two hidden states correspond to the unmethylated (U) and methylated (M) components, with component-specific binomial emission densities. The estimates of the binomial parameters (*p*_*U*_ and *p*_*M*_) and the HMM transition matrix (*i.e*. the collection of probabilities to transition from one hidden state to another) are estimated freely during model training for different sequence contexts, thus requiring no empirical knowledge of the conversion rate. In the present analysis we have used contexts CG, CCG, CWG, CAA, CTA and CCA|CHY (where H= {A,C,T}, W= {A,T} and Y={C,T}), following evidence of their different methylation characteristics [45].

Based on the model fits, the probability that a given cytosine belongs to one of the hidden states is given by the posterior probabilities *γ*_*U*_ and *γ*_*M*_ (Fig. 2d, Methods section). A cytosine’s maximum posterior probability represents its most likely methylation status (Fig. 2d,e), and the magnitude of this probability can be used as a measure of confidence in the underlying status call. In addition to methylation status calls, METHimpute outputs recalibrated methylation levels per cytosine, calculated as *m′* = *p*_*U*_ · *γ*_*U*_ + *p*_*M*_ · *γ*_*M*_ (Fig. 2e). A key feature of METHimpute is its ability to infer the methylation level and status for cytosines with missing data (*i.e*. zero read coverage) or for those with poor read coverage (*i.e*. less than 3 reads). It achieves this inference iteratively during HMM training by borrowing information from neighboring sites. The algorithm therefore outputs complete, base-resolution methylomes, that can otherwise only be obtained through very high-depth sequencing experiments.

**Figure 2.**
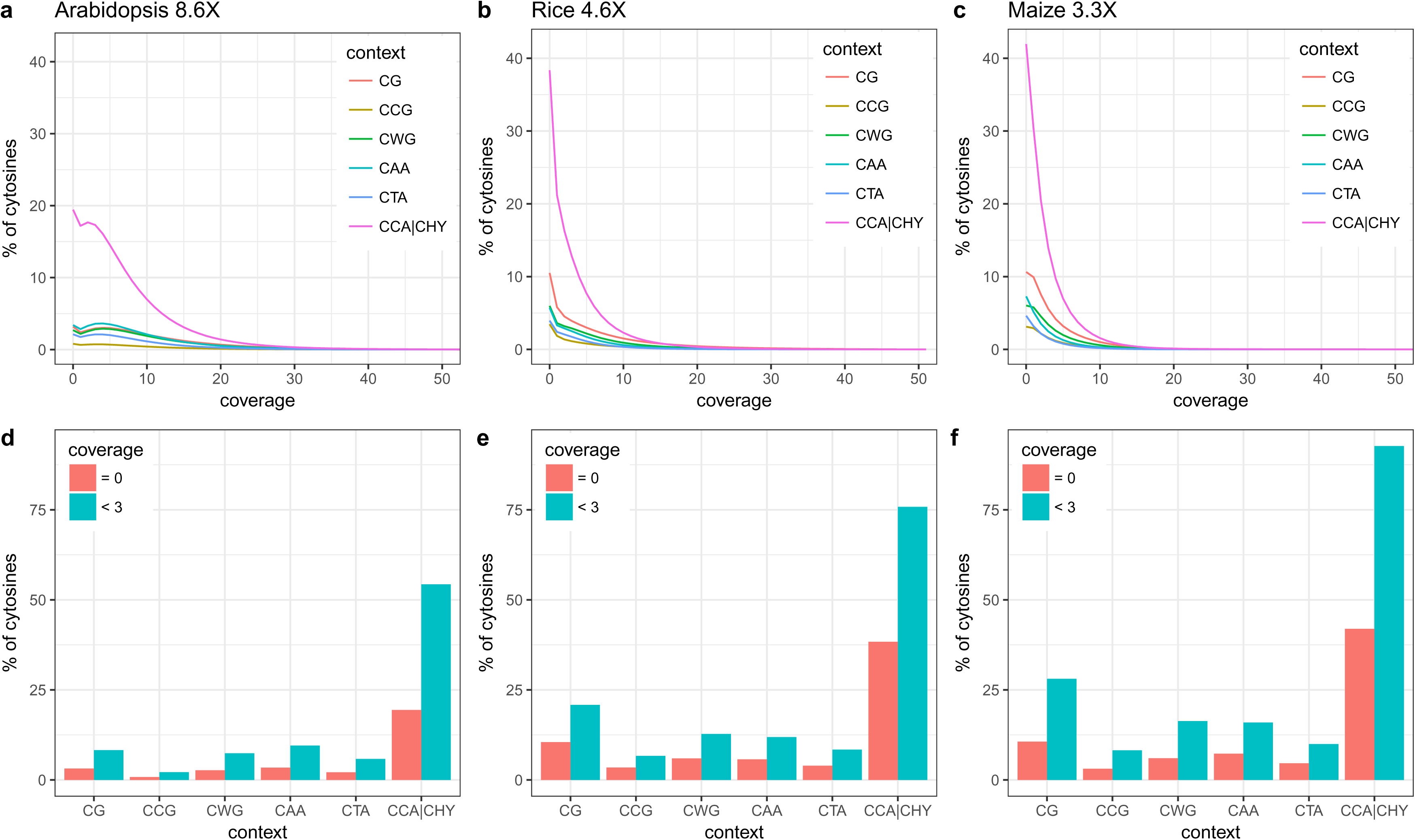
Conceptual overview of METHimpute. **(a)** Cytosines on the sequenced genome are assumed to be either unmethylated or methylated. **(b)** Bisulphite-sequencing and alignment yields methylation levels for each cytosine, *i.e*. the number of reads showing methylation divided by the total number of reads. **(c)** Emission densities for each state are obtained with a binomial test with state-specific parameters. Note that "imputed” cytosines, *i.e*. cytosines without any reads, are treated identically as all other cytosines. However, since the emission densities for all states are 1 for imputed cytosines, the methylation status call is purely driven by the neighborhood of cytosines. **(d)** Model fitting yields posterior probabilities for methylation status calls. **(e)** Inferred methylation status calls and methylation levels.

## Results

Imputation-guided construction of complete Arabidopsis, rice and maize methylomes

To demonstrate the performance of METHimpute we analyzed representative WGBS datasets from *A. thaliana* (Col-0) [41], *O. sativa* (japonica nipponbare) [46], and *Z. mays* (B73) [47]. We chose these three species because they cover a wide spectrum of plant genomes in terms of length and complexity: the *A. thaliana*, *O. sativa* and *Z. mays* genomes are 120 Mb, 374 Mb and 2.1 Gb in length, respectively, and have an estimated repeat content of 10%, 28-35% and 85% [48–51]. The *A. thaliana* data consisted of two replicates (rep.1: 8.6X; rep.2: 15.7X coverage per cytosine per strand), while there were three replicates both for *O. sativa* (rep.1: 7.4X, rep.2.: 6.9X, rep.3: 4.6X) and *Z. mays* (rep.1: 1.6X, rep.2: 3.3X, rep.3: 2.4X). The precise mapping statistics for each dataset are detailed in Table SI-1. Alignment and pre-processing of the data was carried out using a single pipeline as described in the Methods section. Runtimes and memory requirements for METHimpute are listed in Table SI-4.

We examined the genome-wide coverage distributions of each replicate dataset. Despite average coverage being relatively high, a substantial proportion of cytosines had either missing data or low coverage. For instance, in the *A. thaliana* (rep.1: 8.6X), *O. sativa* (rep.3: 4.6X) and *Z. mays* (rep.2: 3.3X) datasets, about 9% (3.71M), 24% (39.54M) and 26% (36.77M) of all cytosines had missing data (*i.e*. zero read coverage) and 24% (10.27M), 49% (79.38M) and 60% (85.5M) were nearly uninformative (here defined as coverage < 3 reads) (Fig. 1d-f and Fig. SI-2 for the other replicates). Interestingly, the genome-wide proportions of missing or uninformative sites were highly context dependent, being highest for CCA|CHY, probably as a result of less unique short read alignments in this context as it is more abundant in repetitive regions of the genome (Fig. SI-3 and Fig. SI-4).

We applied METHimpute to the above-described datasets and evaluated the quality of the resulting methylation calls. For *A. thaliana*, *O. sativa* and *Z. mays*, the algorithm imputed the methylation status of all 3.71M, 39.54M and 36.77M missing data cytosines, respectively, and inferred the methylation status of all 10.27M, 79.38M and 85.5M uninformative cytosines.

### Inferred methylation calls capture known biology

To evaluate the quality of the inferred methylation status calls and levels we examined the per-cytosine posterior probability of being either unmethylated (U) or methylated (M). As mentioned above, this probability represents a measure of statistical confidence in the underlying methylation call, with a value of 1 being the most confident. We found that the distribution of maximum posterior probability values for imputed cytosines shows a clear peak around 1 and a tail of lower confidence values (Fig. 3 and Fig. SI-5 for the other replicates), suggesting that the algorithm produces high-confidence methylation calls for a large proportion of cytosines with missing data. Indeed, 58% (1.50M), 54% (3.96M) and 83% (6.43M) of imputed cytosines in *A. thaliana*, *O. sativa* and *Z. mays* were called with high confidence (defined as posterior probability ≥ 0.9), and these numbers increased to 91% (4.16M), 90% (6.64M) and 93% (9.56M) for cytosines covered by only one or two reads.

**Figure 3.**
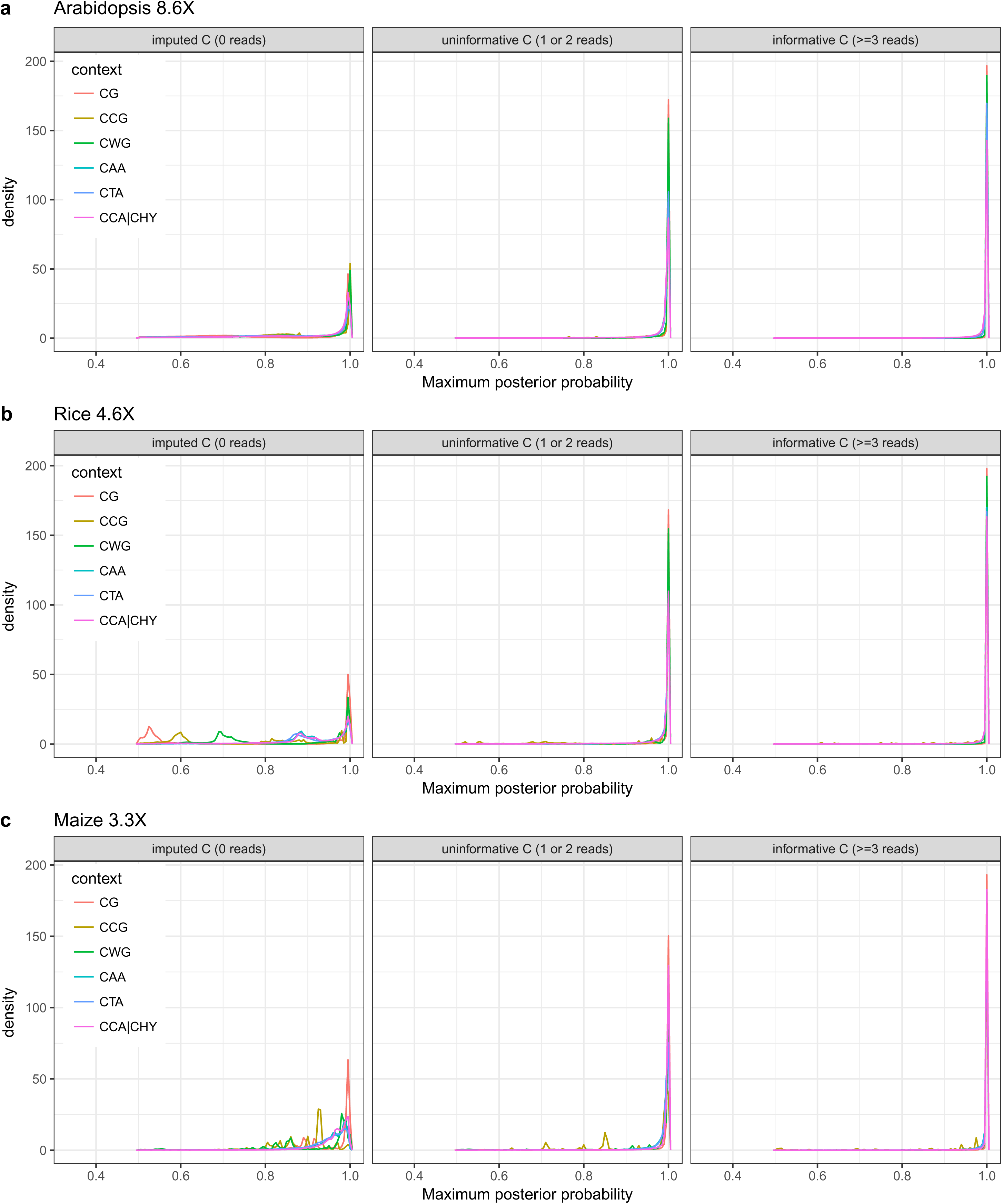
Maximum posterior distributions for imputed cytosines (coverage = 0), uninformative cytosines (coverage = 1 or 2) and informative cytosines (coverage ≥ 3). The figure shows the distributions of the maximum posterior probabilities with density on the y-axis and the maximum posterior probability on x-axis. The maximum posterior probability, *i.e*. the confidence in the methylation status calls, is generally lower for sites with less coverage.

To assess whether the inferred methylation levels are consistent with known biology, we constructed meta-methylation profiles for annotated repeats and genes using cytosines separated in three different categories: informative (coverage ≥ 3), uninformative (coverage = 1 or 2) and imputed cytosines (coverage = 0). Regardless of coverage category, METHimpute confirms that *A. thaliana* TE sequences are heavily methylated in all sequence contexts, with a marked decrease in methylation levels at their 5’ and 3’ ends (Fig. 4b and Fig. SI-6b for the other replicate). The CCA|CHY context shows the lowest methylation levels and CG shows the highest, consistent with [45], and the ordering is conserved for imputed and uninformative cytosines. Similar profiles were detected for repeat elements in *O. sativa* and *Z. mays*, with high CG, CCG and CWG methylation, and very low levels of CAA, CTA, and particularly CCA|CHY methylation, consistent with known results (Fig. 4d,f and Fig. SI-6 for the other replicates) [52].

**Figure 4.**
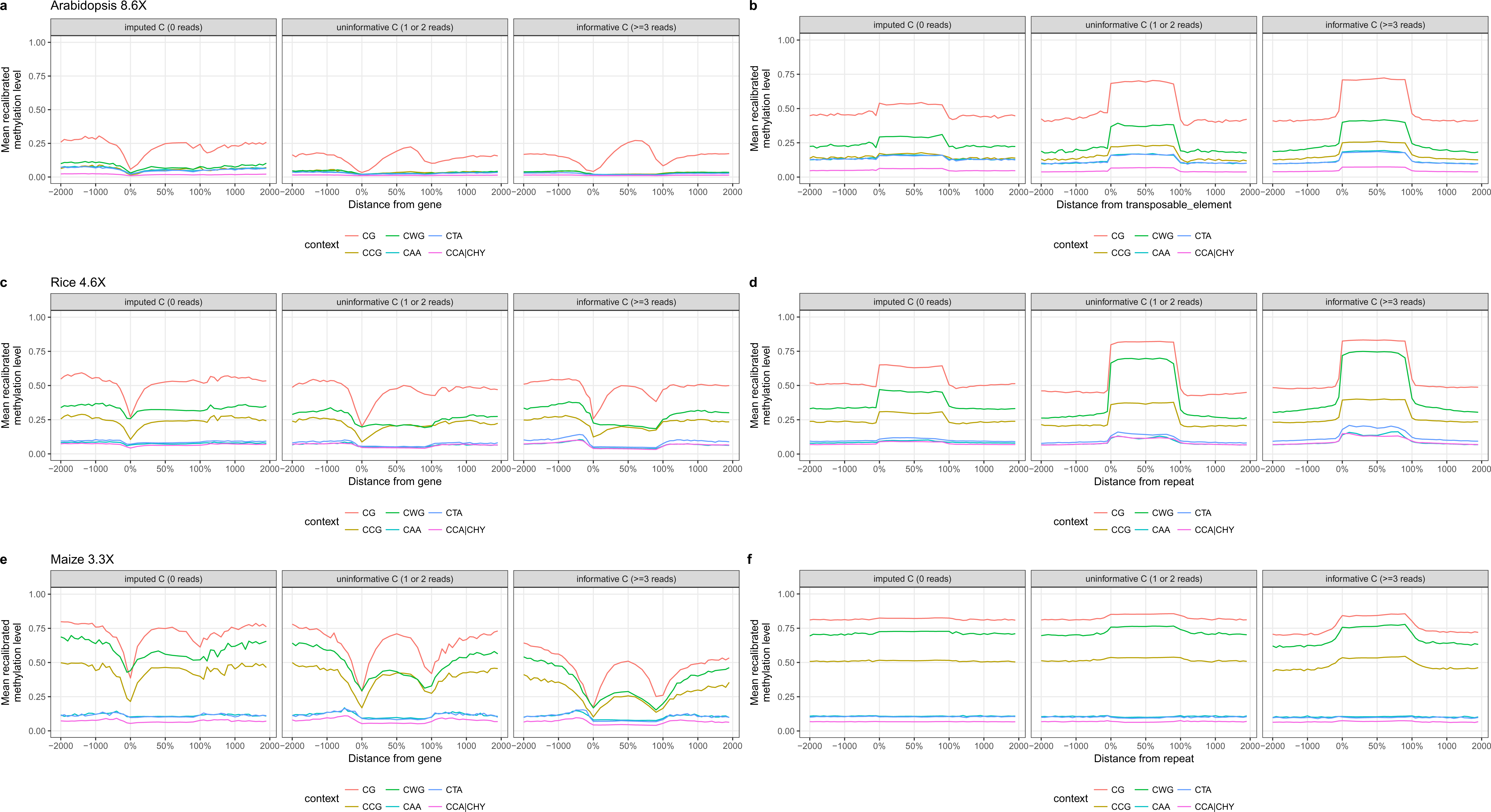
Enrichment profiles for genes (left panels) and transposable elements or repeats (right panels). Sub-panels show the enrichment profiles for imputed (coverage = 0), uninformative (coverage = 1 or 2) and informative cytosines (coverage ≥ 3). See the Methods section for definition of the recalibrated methylation level.

In line with numerous methylome studies in Arabidopsis (e.g. [45,53,54]), ME-THimpute finds that *A. thaliana* genes are intermediately methylated in CG context, and essentially unmethylated at all CHG (CCG, CWG) and CHH (CAA, CTA, CCA|CHY) sites (Fig. 4a and Fig. SI-6a for the other replicate). Genic meta-methylation profiles for *O. sativa* and *Z. mays* were generally similar to those of *A. thaliana* (Fig. 4c,e and Fig. SI-6 for the other replicates), with the exception that both crop species are known to also methylate genic CHG context, probably owing to the fact that genes in these complex genomes often overlap or contain heavily methylated TE or repeat copies.

Taken together the above analyses illustrate two points: first, METHimpute infers annotation-specific methylation profiles that are consistent with published reports; and second, the methylation profiles inferred from imputed or uninformative cytosines recapitulate the patterns seen for highly-informative cytosines, indicating that - regardless of coverage - the inferred methylation calls are robust and biologically meaningful.

### Saturation analysis for the performance assessment of imputed methylomes

METHimpute achieves high quality imputations by leveraging information from neighboring cytosines via the estimated distance-dependent transition probabilities (see Methods section). Therefore, confidence in the imputed calls is higher for cytosines that are closer to informative sites (Fig. SI-7). This spatial dependency remains high over distances of 10-40 bp and then decays to background levels. We reasoned that our imputation method may therefore be relatively robust even in shallow WGBS experiments, as long as enough measured cytosines are available to tag the methylation status of the underlying region.

To test this directly, we implemented a saturation analysis similar to Libertini et al. 2016 [55], where we compared high-coverage datasets with low-coverage subsets of these datasets. Bam files with mapped reads for the Arabidopsis, rice and maize replicates were merged to obtain samples with 23.2X, 18.6X and 7.2X coverage per cytosine per strand, respectively (Table SI-1). These merged files were downsampled to generate a series of reduced datasets, ranging from 90% to 10% of the original data (Table SI-3).

Upon downsampling, the proportion of cytosines with zero read coverage increased from 5% (23.2X) to 31% (13.47M, 2.6X) in *A. thaliana*, from 11% (18.6X) to 40% (65.41M, 1.8X) in *O. sativa* and from 14% (7.2X) to 37% (52.07M, 2.2X) in the *Z. mays* data (Fig. 5d-f). We ran METHimpute on each reduced dataset and calculated the F1-score in the status calls relative to those obtained with the full data. The F1-score is defined as the harmonic mean of precision and recall, and the status calls of the full dataset were assumed as ground truth.

**Figure 5.**
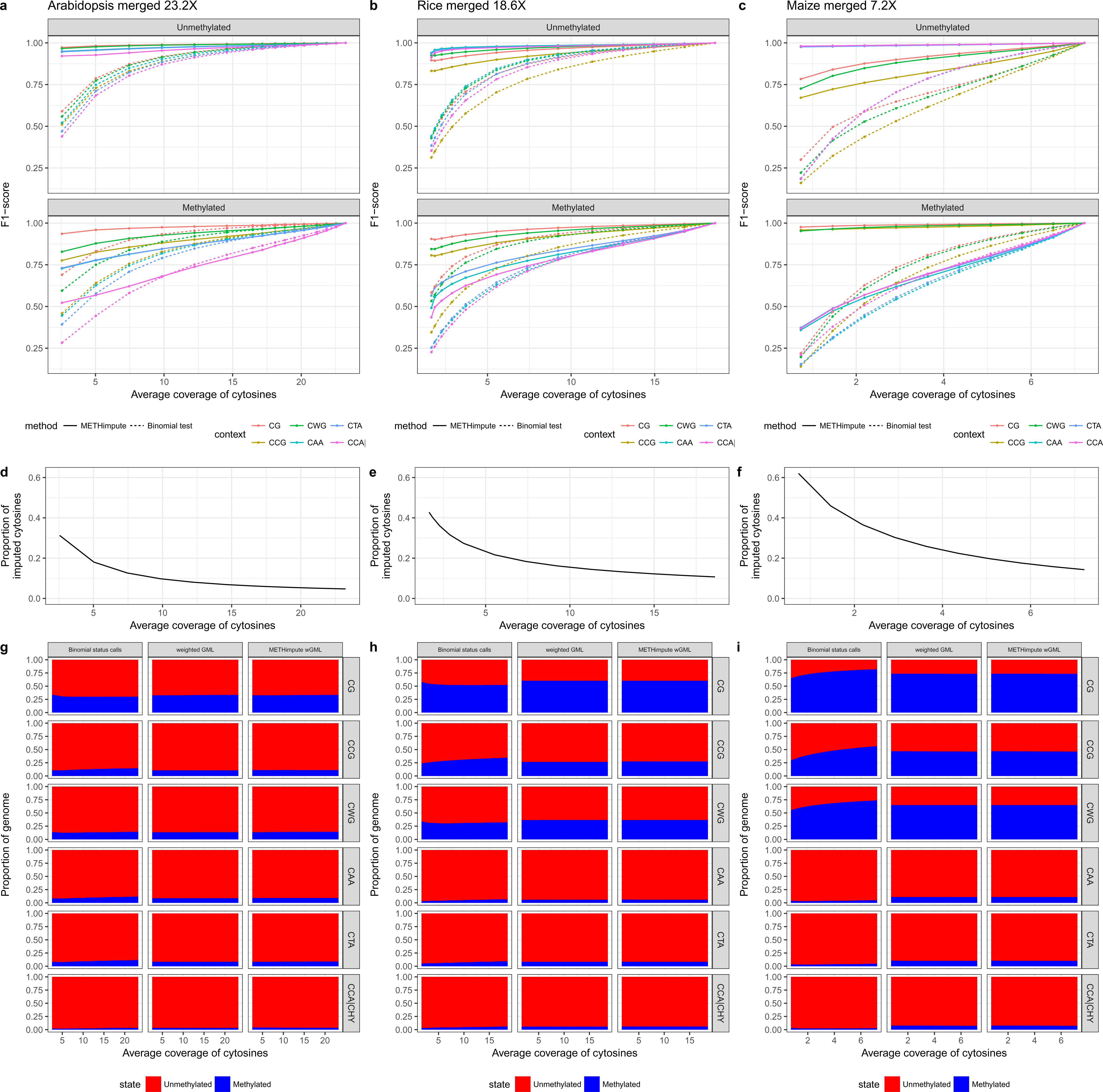
Saturation analysis. **(a-c)** F1-score for METHimpute and the binomial test, compared to the full sample, respectively. The F1-score is the harmonic mean of precision and recall. **(d-f)** Proportion of imputed cytosines. **(g-i)** Proportion of the genome in each state. The x-axes shows the average strand-specific coverage per cytosine.

Our analysis shows that performance remains remarkably high despite drastic decreases in sequencing depth (Fig. 5a-c, Fig. SI-8 with precision and recall, Fig. SI-9 F1-score per context). With data as low as 5X coverage per cytosine (strand-specific), the F1-score was as high as 95% in *A. thaliana* (U: 95%, M: 74%), 97% in *O. sativa* (U: 97%, M: 88%) and 99% in *Z. mays* (U: 99% M: 98%). In general, annotations with a large percentage of missing cytosines in the high coverage datasets were less accurately called upon downsampling (Fig. SI-4). These include in particular transposable elements and repeats. The exception to this trend were 5’ UTRs, which in all three species showed a large percentage of cytosines with missing data but a low amount of miscalled sites upon downsampling.

To put the above accuracy analysis into perspective, we also ran the commonly used binomial testing approach on the reduced datasets. As this method is typically applied to cytosines with sufficient coverage, missing (zero reads) or uninformative sites (< 3 reads) need to be treated as false negatives in the downsampled files. We find that the accuracy obtained with the binomial approach is highly sensitive to average sequencing depth. With only 5X data, the F1-score drops down to 73% (U: 72%, M: 70%) in Arabidopsis, 81% (U: 80%, M: 81%) in rice and 90% (U: 88%, M: 90%) in maize (Fig. 5a-c).

Finally, we also considered the fidelity of the recalibrated methylation levels upon downsampling. Recalibrated methylation levels can be interpreted as the probability of observing a methylated read at a given position, and these recalibrated levels are highly correlated with original methylation levels: For Arabidopsis, rice and maize, the correlation (linear fit) was 0.91, 0.94 and 0.93, respectively (p-value ≤ 2e^‒16^). To assess their fidelity upon downsampling, we calculated the correlation between recalibrated methylation levels per cytosine and per 100bp window to the full coverage dataset, and compared that to the results obtained from the original methylation level (Fig. SI-10). Per-cytosine recalibrated methylation levels show slightly higher correlations than original methylation levels, and with 10% of the original data the correlations for Arabidopsis, rice and maize are 0.89, 0.90 and 0.93, respectively. Window-based recalibrated methylation levels showed the same correlation performance as the original ones, with remarkably high correlations even when only 10% of the original data was retained (0.95, 0.95, 0.83 for Arabidopsis, rice, maize). These results suggest that recalibrated methylation levels can be used for downstream methylation analysis, since they are correlated to original methylation levels and are robust upon downsampling, while providing cytosine-level information even at low sequencing depth.

Overall, both for status calls and for recalibrated methylation levels, METHimpute produces robust results even at very low sequencing depth, suggesting that the algorithm offers a cost-effective solution for methylome studies of large genomes and for population-level studies involving a large number of samples.

### Re-calibrated estimates of genome-wide and context-specific methylation levels

Plant species differ greatly in their genome-wide methylation levels (GMLs, *i.e*. the proportion of cytosines that are methylated) [3,4]. In a recent survey of about 30 angiosperms, GMLs were found to be as low as 5% in *Theobroma cacao* to as high as 43% in *Beta vulgaris*, with a mean of about 16% [3,39]. Much of this diversity appears to be the result of differences in genome size and repeat content, as well as differences in the efficiency of DNA methylation maintenance pathways [28]. Precise estimates of GMLs are important for studying the evolutionary forces that shape plant methylomes over short and long time-scales, and for understanding genome-epigenome co-evolution. However, obtaining GML estimates based on WGBS data is not trivial, as they are highly dependent on the method used for methylation status calling and on the depth of the sequencing experiment. In *A. thaliana*, for instance, reported GML estimates vary widely between studies. This dependency is even larger when considering context-specific GMLs (*i.e*. the proportion of methylated cytosines in a given context; CG-GMLs, CHG-GMLs, CHH-GMLs), with CHH-GMLs being by far the most variable between studies, with reported values ranging from as low as 1.51% [1] to as high as 3.91% [3].

In order to bypass many of the statistical issues in calling methylation states, especially in shallow WGBS data, recent studies have proposed so-called weighted genome-wide methylation levels (wGMLs) as a proxy for GMLs. A wGML is a non-statistical measure which is obtained by counting the number of methylated reads over the total number of reads at the genome-wide scale. Fig. 5g-i shows clearly that wGMLs are robust upon down-sampling in any sequence context in the *A. thaliana*, *O. sativa* and *Z. mays* data, thus justifying its use. By contrast, GMLs calculated from cytosine-level binomial status calls (i.e. #mC/all Cs) are highly unstable, particularly in non-CG contexts and when sequencing depth is low.

In order to assess whether the re-calibrated methylation levels provided by METHimpute can also be used to obtain robust estimates of GMLs, we calculated wGMLs by summing the per-cytosine re-calibrated methylation level genome-wide, weighted by coverage. Using this measure we find that METHimpute-derived wGMLs perform nearly identical to naive wGMLs, both in terms of robustness and magnitude (Fig. 5g-i, Fig. SI-11 with replicates). This demonstrates that METHimpute recalibrated levels are consistent with original methylation levels and known biology not only at the individual cytosine level, but also aggregated over 100bp windows and genome-wide, with the added advantage that they are available for all positions in the genome.

### METHimpute facilitates insights into bisulfite conversion rates

One source of measurement noise in WGBS data is the bisulfite conversion procedure prior to sequencing. Bisulfite treatment of DNA is typically performed long enough so that all unmethylated cytosines are converted to uracils. The conversion success (or rate) is typically high. Most studies report conversion rates of about 0.99, implying that only about 1% of all unmethylated cytosines failed to convert. Knowledge of this rate is important not only to verify that bisulfite reaction was efficient but also to be able to separate biological signal from noise in downstream analyses of the data. Empirical estimates of the conversion rate are often obtained by including unmethylated chloroplast and virus genomes as controls in the WGBS workflow, and counting the number of non-converted cytosines from the mapped reads.

A helpful byproduct of the METHimpute fitting procedure is that the conversion rate can be directly estimated from the sequenced material without requiring auxiliary information from chloroplast or virus genomes. METHimpute achieves this in the HMM framework by estimating the probability, *p*_*U*_, of finding a methylated read given that the underlying cytosine is unmethylated (see Methods), which can be used to derive the conversion rate. To obtain these rates we focus on estimates of *p*_*U*_ in context CG to exclude potential biases arising from the “fuzzy” maintenance of methylation at CHG and CHH sites. For *A. thaliana* and *Z. mays* our estimated conversion rates were 0.989 and 0.961, respectively, which is remarkably close to chloroplast-based estimates of 0.993 and 0.970.

Although bisulfite conversion kits and protocols have been optimized to achieve the highest conversion rate possible the specificity of the reaction is not perfect. A well-known trade-off is that some methylated cytosines can be accidentally converted to uracils, and are later falsely detected as unmethylated. Some controls (commercial or artificially methylated DNA fragments) are available to estimate this inappropriate conversion rate, but, to our knowledge, they are not systematically used in WGBS experiments. Some studies using such controls have shown that the inappropriate conversion rate (% of methylated cytosines converted to uracils) ranges from 0.09% to 6.1% depending on the kit and protocol used [56–58].

METHimpute approximates this value by estimating the parameter *p*_*M*_ for the *M* component (see Methods), which can be used to calculate the probability of finding an unmethylated cytosine given that the underlying cytosine is truly methylated. Again, focusing on CG sites, we estimate the methylated cytosines conversion rate at 6.3%, 11.5% and 16% in *O. sativa*, *Z. mays* and *A. thaliana*, respectively. Although these estimates are close to the empirical rates reported in the literature, they are slightly biased upward most likely owing to the fact that the parameter *p*_*M*_ is partly confounded with methylation variation arising from cellular heterogenity in the sampled tissues. We therefore suspect that our estimates become more accurate in situations where tissue heterogeneity is minimized.

Nonetheless, the ability of METHimpute to provide an accurate estimate of the conversion rate for unmethylated cytosines and an upper-bound estimate for methylated cytosines could be utilized to calibrate WGBS experiments in the laboratory when no controls are available.

## Discussion

A key advantage of WGBS over alternative measurement technologies is its ability to provide cytosine-level measurements from bulk and - more recently - also from single cell data. Since its first application in the model plant *A. thaliana* in 2008 [53,54], WGBS has become an integral tool for studying the methylomes of increasingly large plant genomes and for surveying patterns of natural methylome variation within and among plant species. However, the relatively high costs associated with this technology pose limits on the sequencing depths that can be achieved within most experimental budgets. A typical solution is to sequence methylomes far below saturation, which results in substantial measurement noise and missing data at the level of individual cytosines.

Here we introduced METHimpute, an imputation-based HMM for the construction of complete methylomes from shallow or deep WGBS data. Our analyses showed that the algorithm can impute the methylation status of cytosines with missing data (*i.e*. zero read coverage) or uninformative coverage (*i.e*. coverage of less than 3 reads), as well as their recalibrated methylation levels. We demonstrated that these imputations are not only statistically robust, but also biologically meaningful. Our estimates suggest that routine use of this algorithm could reduce sequencing costs of typically sized methylome experiments by about 50% without a substantial loss of biological information. The method works with small, streamlined genomes like that of Arabidopsis but also with large, repeat-rich genomes like those of most commercial crops, thus making it a flexible software tool for the analysis of DNA methylomes of a wide spectrum of species.

We recommend the use of METHimpute instead of the binomial test for the analysis of WGBS data whenever methylation status calls are required. Furthermore, METHimpute solves the problem of missing data in population epigenetic studies, which will facilitate the estimation of epigenetic mutation rates and methylation site frequency spectrum analyses.

METHimpute is implemented as an R-package and seamlessly integrates with the extensive bioinformatic tool sets available through Bioconductor. The algorithm has been extensively tested in plants, but it should also be applicable in non-plant species.

## Methods

### Hidden Markov Model for methylation calling

#### Outline

We define an *N* = 2 state Hidden Markov Model (HMM), where the states *i* represent unmethylated (U) and methylated (M) cytosines. The emission densities for each state are binomial distributions, which can be interpreted as a binomial test on the number of methylated counts *m* over total counts *r*. The probability parameter *P*_*i*_ of the binomial test can be interpreted as the probability of finding *m* methylated counts out of *r* total counts, given the state *i*. Note that in this definition 1 — *p*_*U*_ is the conversion rate, *i.e*. the probability of a read showing non-methylation when the cytosine is indeed non-methylated. Cytosines are not equally spaced in the genome, and we therefore chose a distance dependent transition matrix **A**, where the distance dependent change in transition probabilities is modeled by an exponential function. Furthermore, to account for different sequence contexts, we implemented context-specificity for both the binomial test and the transition probabilities.

### Mathematical description

The probability *P* of observing methylated *m*_*t*_ and total *r*_*t*_ read count at a particular cytosine *t* in context *c*_*t*_ can be written as

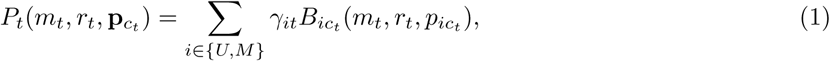

where *γ*_*i*_ are the posteriors (mixing weights) and *B*_*i*_ are binomial distributions with context-specific parameter *p*_*ic*_. The binomial distribution is defined as

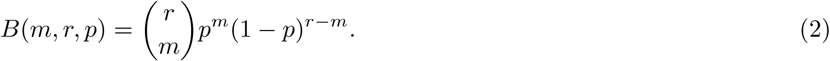

All probability parameters of the binomial tests (*i.e*. the probabilities of a success) are estimated freely during model training (next section). For *C* = 6 contexts and *N* = 2 states, *N · C* =12 independent parameters *p*_*ic*_ need to be fitted.

The distance dependent transition probabilities from cytosine *t* in state *i* to cytosine *t* + 1 in state *j*, separated by distance *d*_*t*,*t*+1_ and in transition context *c*_*t*,*t*+1_, can be described as

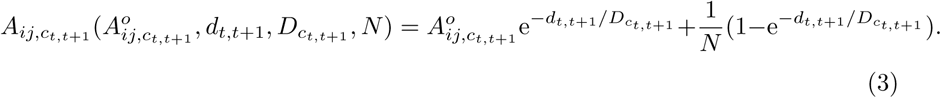

Here, 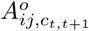 are the transition probabilities without distance dependency (or for adjacent cytosines with *d*_*t*,*t*+1_ = 0). *D*_*c*_*t*,*t*+1__ is a constant that reflects how fast neighboring cytosines lose correlation. The distance dependency is constructed in such a way that all transitions *A*_*ij*,*c*_*t*,*t*+1__ are equally likely for an infinite distance *d*_*t*,*t*+1_ = γ Note that for *C* =6 contexts the model has *C · C* = 36 transition contexts and thus 36 different transition matrices with dimensions *N* × *N*.

The constants *D*_*c*_ are determined by a non-linear least-squares (nls) fit to the correlation decay between cytosines in transition context *c*_*t*,*t*+1_ (see Fig. SI-12 for all used transition contexts). The formula for the fit is *y*_*c*_(*d*) = *a*0 * *e*^‒*d/D*_*c*_^, where *y*_*c*_ is the correlation between neighboring cytosines at distance *d* in transition context *c*. The parameters *a*0 and *D*_*c*_ are fitted by the nls-fit.

An important point is that the correlation is calculated between adjacent cytosines, with no other cytosines in between. This reflects the definition of the transition probabilities in the Hidden Markov Model, where transitions are defined from one cytosine to the next in the sequence.

### Model fitting

Model parameters are fitted with the Baum-Welch algorithm [59]. The distance-dependent transition probabilities require modified updating formulas compared to a standard Baum-Welch algorithm without distance dependency. The derivation of the modified updating formulas is detailed below, and uses notation introduced in [60].

The conditional expectation *Q* that needs to be maximized can be written as

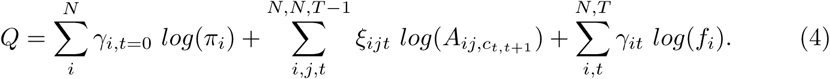

The updated transition probabilities 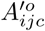 can be obtained by solving 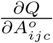 using the method of Lagrange multipliers to deal with the constraint 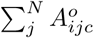

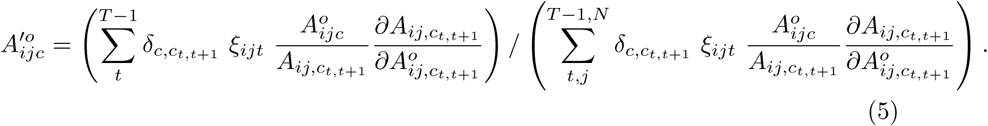

Here, *δ*_*c*,*c*_*t*, *t*+1__ is the Kronecker delta function, which ensures that only terms in the correct transition context *c* are included into the sum.

Similarly, the updated parameters for the binomial test can be obtained by solving 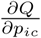. For independent binomial tests, this yields

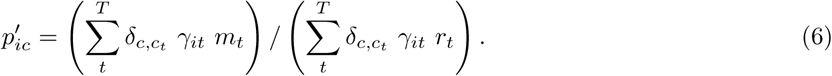

The methylation status *i*_*t*_ is determined by maximizing over the posterior probabilities *i*_*t*_ = *argmax*_*i*_(*γ*_*it*_).

Finally, we can use the posterior probabilities *γ*_*U*|*M*,*t*_ and estimated parameters *p*_*ic*_ to define a recalibrated methylation level 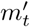 that is defined on every cytosine *t* in the genome and can serve as input for other applications:

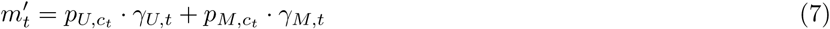

### Plants DNA methylation data

In this study, we used published data (fastq files containing bisulfite sequenced reads) from three model plant species to test METHimpute: *Arabidopsis thaliana*, rice (*Oryza sativa* Japonica cv. Nipponbare) and maize (*Zea mays* B73). We used three replicates for rice and maize, and two replicates for *Arabidopsis*. Each sample was mapped to the latest available version of the reference genome for this species. Details and references on these datasets, reference genomes and annotations files, as well as additional alignment metrics can be accessed in Table SI-2.

### Mapping of bisulphite sequenced (BS-seq) reads and construction of DNA methylomes

Read sequences (Table SI-2) were quality trimmed and adapter sequences were removed with Cutadapt (version 1.9; python version 2.7.9; [61]). Trimming was performed on both ends using the single-end mode and the quality threshold was set to a phred score of 20 (q = 20). We applied the default error rate of 10% for the removal of the adapter sequences. Afterwards, we discarded reads shorter than 40 base pairs. Reads were subsequently mapped to an indexed genome. The maximum allowed proportion of mismatches was set to 0.05 (m = 0.05, 5 mismatches per 100 bp) and the maximum insert size was set to 1000 bp (X = 1000). BS-Seeker2 (v2.0.10; [44]) using Bowtie2 (version 2.2.2; [62]) was chosen for the alignment of the reads. Samtools (version 1.3.1; using htslib 1.2.1; [63]) was used to remove duplicates (samtools rmdups) and to sort bam files (samtools sort). Methylomes were subsequently constructed through the bs_seeker2-call_methylation.py module from BS-Seeker2 (v2.0.10; [44]). CGmap files containing methylome information were used as an input for METHimpute.

## Availability of data and materials

METHimpute can be downloaded from https://github.com/ataudt/methimpute.

## Author′s contributions

A.T., M.C-T. and F.J. conceived this project; A.T. implemented the model with input from M.C-T, F.J. and D.R.; A.T., D.R., R.W. and A.V. analyzed the data; A.T., M.C-T. and F.J. wrote the paper.

## Acknowledgments

We thank R. Schmitz and C. Niederhuth with their help in accessing the maize and rice annotation files and providing data, and N. Springer and R. Schmitz for their quick feedback on this project. FJ and DR acknowledge support from the Technical University of Munich-Institute for Advanced Study funded by the German Excellence Initiative and the European Union Seventh Framework Programme under grant agreement #291763. MCT acknowledges support from the Helmholtz Association’s Initiative and Networking Fund and from the University of Groningen (Rosalind Franklin Fellowship).

